# RBC Deformability During Acute Lymphoblastic Leukemia Treatment

**DOI:** 10.64898/2026.10.29.762767

**Authors:** Tuna Pesen, Yigitcan Koc, Beyza Berin Bagislar, Nurcan Alhan, Bora Akgun

## Abstract

Philadelphia chromosome positive acute lymphoblastic leukemia (Ph+ ALL) is the most common genetic subgroup in adult ALL, and its incidence increases with age. In this longitudinal single-subject study, we investigate erythrocyte deformability as a disease-associated biomechanical marker in parallel with clinical, hematological, and molecular response throughout the treatment process in a 75-year-old patient diagnosed with Ph+ ALL. Using optical tweezers, we performed high-resolution single-cell mechanical stretching experiments to quantify the deformability of individual RBCs obtained from the patient and a healthy control. Our measurements reveal a pronounced relation between state of diseases/therapy and RBC deformability in the ALL patient, indicating systemic alterations in cellular biomechanics associated with leukemic pathophysiology. These findings suggest that changes in RBC mechanical properties, probed through non-invasive optical tweezers-based techniques, may accompany disease presence and progression.

## Introduction

Acute lymphoblastic leukemia (ALL) is an aggressive hematologic malignancy characterized by the uncontrolled proliferation of lymphoid precursor cells. In adult ALL, the most common genetic subgroup is Ph+ ALL, whose incidence increases with age. In this disease, formerly associated with poor prognosis, clinical outcomes have significantly improved and long-term survival rates have now risen to approximately 75–80% with the introduction of TKI-based therapies and immunotherapies.^1,2^ In patients over 65 years of age with Ph(+) ALL, the current approach consists of regimens including low-dose chemotherapy and TKI, tailored according to the patient’s age and performance status.^3–5^

The pathophysiology of ALL involves remodeling of the bone marrow microenvironment, which disrupts normal hematopoiesis.^6,7^ Infiltration of the bone marrow by leukemic cells suppresses healthy blood cell production, leading to cytopenias such as anemia, neutropenia, and thrombocytopenia.^8^ Among these manifestations, anemia is one of the most common clinical findings and has traditionally been attributed to insufficient erythropoiesis caused by leukemic infiltration.^9,10^ However, the mechanisms contributing to anemia may not be solely limited to a reduction in red blood cell (RBC) number.

The functional integrity of RBCs, particularly their deformability, defined as the ability to undergo shape changes under applied mechanical forces, is essential for efficient oxygen transport to tissues^11–13^ and for maintaining healthy microcirculation.^14^ Because RBC deformability is a complex biomechanical property determined by membrane elasticity, cytoplasmic viscosity, cell geometry, and the applied force field, numerous experimental approaches have been developed to quantify it under conditions that mimic macro- and microcirculatory stresses.^15^ Deformability also plays a crucial role in splenic filtration, where mechanically impaired RBCs are retained within splenic slits and removed through a natural quality-control mechanism.^16^ In various pathological conditions, oxidative stress, inflammatory cytokines, and metabolic alterations can damage the erythrocyte membrane and cytoplasm, ultimately impairing deformability.^17–19^

Leukemia-associated systemic inflammation and oxidative stress are thought to affect erythrocyte structure and function.^20,21^ Indeed, a variety of RBC membrane abnormalities, including pore formation, loss of the characteristic biconcave morphology, and irregular surface protrusions, have been reported in leukemic patients.^22^ Ultrastructural analyses have further revealed nanoscale pores on the RBC membrane surface, suggesting substantial impairment of RBC structural integrity.^22^ Consistent with these findings, altered blood rheology has also been described in leukemia. Despite comparable erythrocyte levels, leukemic patients exhibit a markedly reduced hemoglobin-to-viscosity ratio due to increased blood viscosity and leukocrit, resulting in compromised microcirculatory flow and tissue oxygen delivery.^23^

Despite these observations, changes in RBC deformability specifically in ALL remain poorly understood, particularly in elderly patients, who represent a clinically vulnerable population with unique therapeutic challenges. In literature, the direct measurements of RBC deformability in elderly patients with ALL are scarce.

Therefore, the aim of this study was to measure erythrocyte deformability in a 75-year-old patient diagnosed with ALL and to discuss the findings in the context of anemia and disease pathophysiology. This study seeks to provide preliminary evidence supporting the hypothesis that erythrocyte dysfunction may contribute to anemia in ALL and to establish a basis for future comprehensive investigations.

## Case Presentation and Treatment

A 75-year-old man with no known systemic diseases other than hypertension presented in May 2025 with complaints of gradually increasing fatigue, malaise, and widespread body pain for about two weeks. No significant pathological findings were detected on physical examination. Anemia (HB: 9.2 g/dl) and thrombocytopenia (PLT: 35,000/µL) were detected in laboratory examinations performed at admission. Viral serology, rheumatologic serology, and nutritional studies aimed at determining the etiology were normal, and no pathological findings were found on Thorax and Whole Abdomen CT performed for malignancy exclusion. Due to the sudden onset of anemia and thrombocytopenia, bone marrow aspiration and biopsy were performed. In the bone marrow smear, lymphoblasts were observed at approximately 80%. Flow cytometry analysis was consistent with B-lymphoblastic leukemia. The bone marrow biopsy was reported as approximately 95% lymphoblastic cell infiltration in a hypercellular bone marrow background. In cytogenetic and molecular studies, a 46 XY karyotype and t(9-22) translocation were found positive, and the BCR-ABL1 p190 transcript level was detected as 19% by PCR. Evaluation of cerebrospinal fluid (CSF) performed at diagnosis revealed no central nervous system (CNS) involvement.

A low-intensity chemotherapy regimen (cyclophosphamide, vincristine, and dexamethasone) was initiated in the elderly and frail patient. On day 15 of treatment, 140 mg dasatinib daily was added to the regimen upon detection of 19% positivity for the BCR-ABL1 p190 transcript in molecular analyses. Treatment was then continued with dasatinib, dexamethasone, and vincristine. For central nervous system prophylaxis, the patient received intrathecal therapy (MTX-ARA-C-dexamethasone) four times. No malignant cells were observed in the cerebrospinal fluid cytologies monitored.

BCR-ABL P190 PCR negativity was achieved in the third month of treatment. The patient’s treatment was continued with Dasatinib at 140 mg per day, as BCR-ABL P190 PCR negativity persisted during follow-up. In the ninth month of treatment, the patient developed secondary grade 1 pleural effusion due to dasatinib. Consequently, dasatinib treatment was temporarily interrupted and the patient was treated with diuretics and dexamethasone. After regression of the pleural effusion and achievement of clinical stabilization, Dasatinib was restarted at a dose of 70 mg/day. During follow-up, the patient’s pleural effusion did not recur and monitoring continues uneventfully with BCR-ABL P190 PCR (-).

In the pre-treatment period, the patient had significant anemia (Hb: *≈* 9-10 g/dL), and erythrocyte deformability analyses also indicated an initial impairment. A response in peripheral blood counts was observed early during the first treatment cycle with Dasatinib. Concurrently, and correlating with the clinical response, repeat erythrocyte deformability analyses performed on post-treatment blood samples revealed a marked improvement compared to the baseline values. This finding suggests that hematological remission was paralleled not only by an increase in cellular counts but also by an improvement in the functional properties of RBCs. Throughout the treatment course, the patient tolerated Dasatinib well overall.

## Methods

### Optical Tweezers Analysis

Optical tweezers operate on the principle of momentum transfer, where forces exerted by a strongly focused laser beam capture dielectric microparticles or biological cells.^24^ For a trapped cell, the equilibrium between scattering and gradient forces ensures stable confinement and precise spatial manipulation.^25^ In stretching-based experiments, cells are subjected to controlled optical loads; the resulting mechanical deformation is subsequently quantified through the analysis of high-resolution microscopy images. By utilizing highly focused dual-beam configurations, this technique enables the investigation of cell mechanics at the single-cell level with high sensitivity.^26^

In the present study, dual-beam optical tweezers were employed to characterize the mechanical behavior of red blood cells (RBCs) by monitoring their deformation under calibrated optical forces. The degree of morphological change induced by these forces serves as a robust indicator of cellular deformability. Specifically, this research evaluates whether leukemia-related physiological alterations correlate with detectable changes in the biomechanical response of RBCs. A schematic overview of the integrated experimental and analytical workflow is illustrated in Fig. 1.

**Figure 1.**
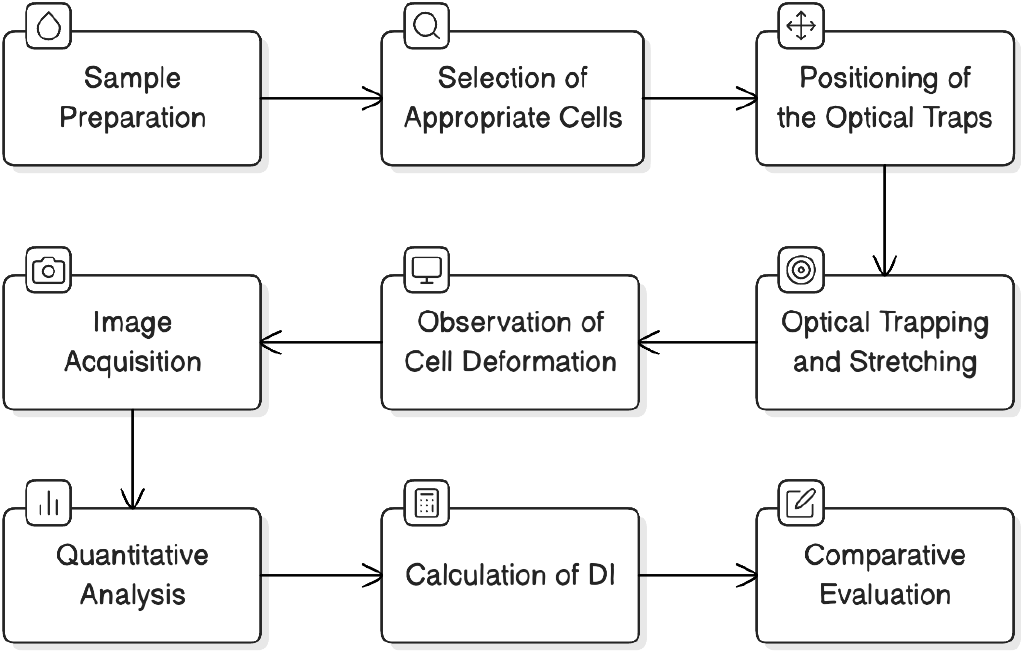
Schematic workflow diagram of the optical tweezers analyses used for single-cell manipulation and deformability measurements.

Across the treatment course, a total of 494 individual red blood cells from the patient were trapped, stretched and analyzed, distributed over eight sampling time points as follows: day 17, n = 31; day 20, n = 42; day 31, n = 83; day 48, n = 88; day 88, n = 60; day 96, n = 69; day 102, n = 60; and day 123, n = 61 (days counted from 16 May 2025). From a healthy donor, 38 cells were measured under identical conditions, served as the healthy reference, giving 532 analyzed cells in total. Each cell was stretched once, so the number of cells defines the unit of replication within a time point, whereas the patient constitutes a single subject followed longitudinally.

### Experimental Setup

All stretching experiments were performed using a commercial optical tweezers system (Zeiss PALM Micro Tweezers). The setup was built around an inverted microscope integrated with a 1064 nm laser source, beam-shaping optics, a high-numerical-aperture objective, a sample stage, and an imaging unit. For optical trapping, an EC “Plan-Neofluar” 100×/1.3 Oil Iris M27 objective lens was used, with a numerical aperture of 1.3 and an approximate transmittance of 50% at the laser wavelength. Although the laser power behind the objective was 800 mW, the effective power at the sample back focal plane was calculated to be 200 mW for each optical trap due to the objective transmittance.

The focused laser beam generated stable optical traps that enabled the manipulation and deformation of individual cells. During the experiments, the hanging-drop slide containing the cell suspension was positioned on the microscope stage, and single RBCs were selected under visual guidance for trapping. Images of the cells were recorded throughout the trapping and deformation process for subsequent analysis. The general optical trapping and stretching configuration used in this study is illustrated in Fig.2.

**Figure 2.**
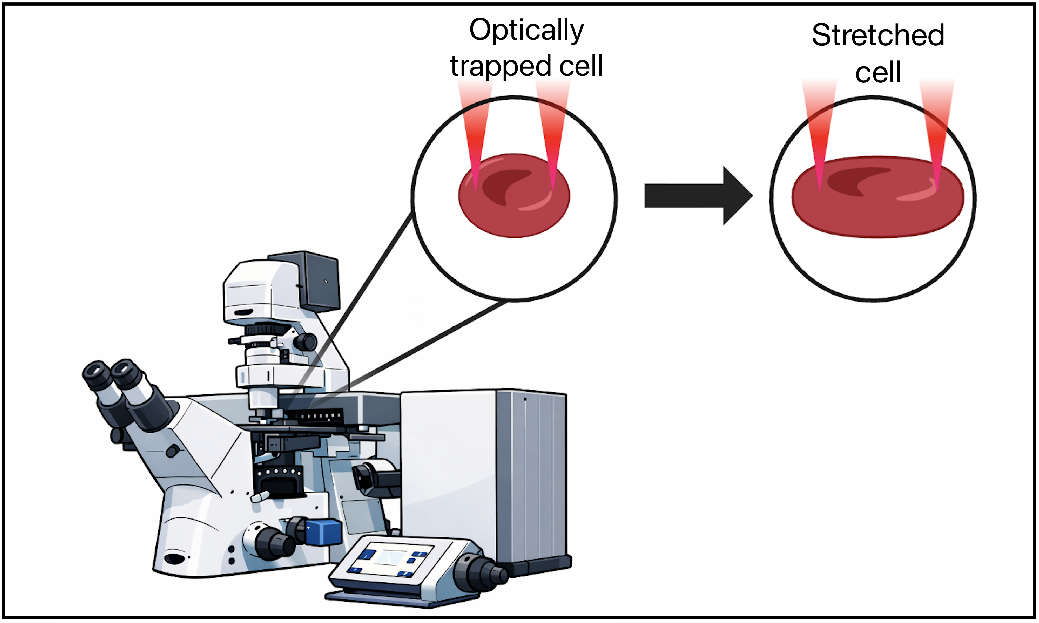
Illustration of the optical trapping and stretching principle. A red blood cell is optically manipulated and deformed under controlled laser-traps.

### Collection of blood samples

A 75-year-old ALL patient and 35-year-old one healthy woman volunteered for this study and written informed consent was obtained from all the participants. The experimental protocols, including blood sample, were approved by Boğaziçi University Science and Engineering Fields Human Research Ethics Committee (FMINAREK) with the ethical permission number 2025-15. Blood samples were taken using a needle from the participants’ veins. All methods were performed in accordance with Declaration of Helsinki.

### Sample Preparation

For deformability measurements, 0.1 µL of blood taken from the original hemogram tubes that were taken in the hospital was diluted in 1 mL of phosphate-buffered saline containing 100 µL bovine serum albumin to obtain a suspension suitable for single-cell optical stretching experiments. From this mixture, 70 µL was placed on a microscope slide and covered with a glass that is treated with BSA. The cover glass was prepared by applying 15 µL BSA and drying it in an incubator at 40^*°*^C for 10 min before use. The edges of the cover glass were sealed with nail polish. The sample preparation procedure and the overall optical tweezers setup were largely similar with the former study.^27^ This treatment was introduced to reduce the adhesion effects of RBCs to the glass surface during measurements. In addition, all experiments were performed at room temperature to maintain consistent handling conditions throughout the study.

### Cell Manipulation and Stretching Protocol

Selected single RBCs were manipulated using the same dual-beam optical tweezers configuration described above. Prior to stretching, the traps were aligned near the opposite ends of the selected cell with an initial separation of 5 *µ*m while the laser was off. After alignment, the laser was switched on and one trap was moved at a constant velocity of 1 *µ*m/s for 10 s, while the other remained stationary. This resulted in controlled stretching of the cell along its axial direction. The cell elongation was monitored throughout the stretching process until maximum deformation was reached, followed by relaxation after unloading or escape. Representative pre-stretch and stretched cell images are presented in Fig.3, and the recorded image sequences were used for subsequent deformability analysis. Using the equation Eq.1, deformability index was calculated for each cells using the analysis method given in the former study.^27^

**Figure 3.**
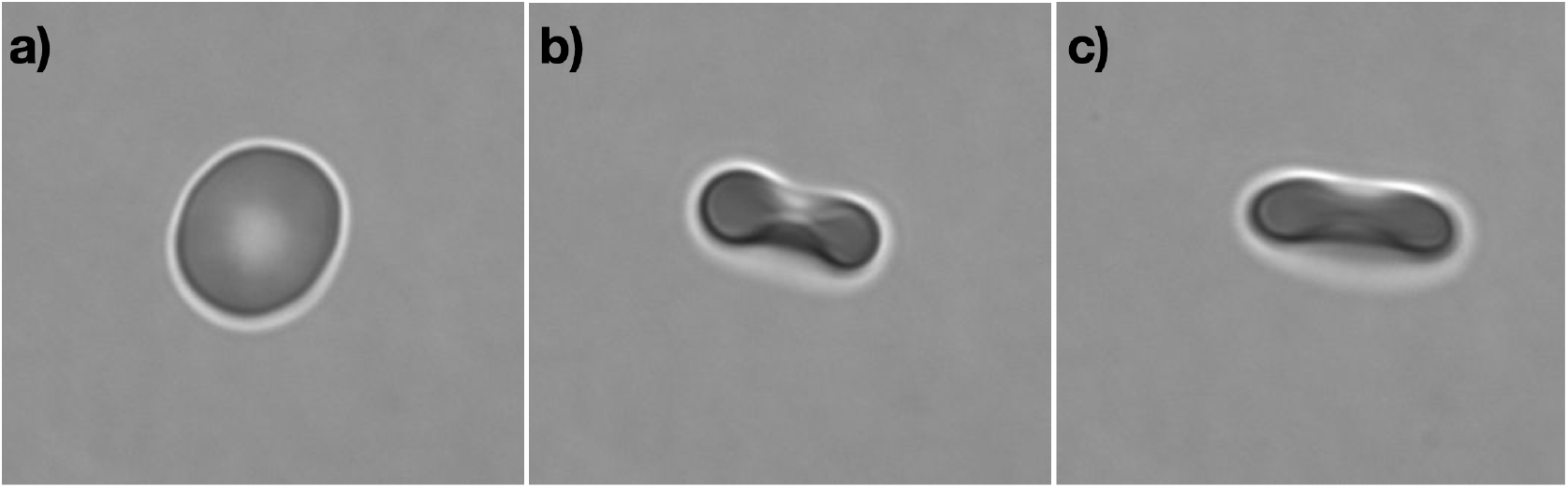
Representative images showing the deformation of an RBC: a) before trapping, b) upon capture by the optical tweezers, and c) after stretching.

### Quantification of Deformability

The deformability of the RBC’s was quantified by utilizing the recorded microscopy images during the stretching procedure. These image-based parameters were deduced and analyzed to determine the cell length before, during, and after the application of optical force. The deformation response of each cell was evaluated by comparing its initial morphology to its shape under trapping force. A deformability-related metric which is deformability index (DI), was defined based on the relative change in cell dimensions during stretching, as follows:

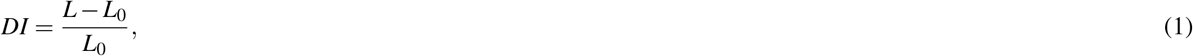

where *L*_0_ denotes the initial axial length of the cell and *L* represents the maximum stretched length captured during the experiment. The observed deformation patterns were interpreted in relation to membrane elasticity and overall mechanical integrity of RBCs to identify possible deviations associated with leukemia-related alterations.

## Results

The deformability of RBCs was measured for 8 hemogram tube samples taken at different time points during the treatment. The initial 4 measurements were made while the patient was still in the chemotherapy period (the region highlighted with a green line in Fig.4). The results showed relatively low mean deformability index values in the range of 0.11–0.13. The rest of the measurements were made after the patient had already stopped chemotherapy treatment. The fifth measurement indicated an elevated DI score of 0.16. Since the patient’s medical condition was not satisfactory at that point, it was decided that a temporary interruption of dasatinib treatment might be beneficial. Evaluation of the sixth and seventh samples demonstrated even higher DI values in the range of 0.22–0.24. After that, the patient continued dasatinib treatment, and the last measurement showed a value within the expected range, as the patient was showing strong indications of remission.

In order to provide a healthy reference for the interpretation of patient-related DI measurements, the DI distribution of the control group was evaluated. The results of the control group are also summarized in Fig. 4. The rightmost boxplot presents the overall distribution of control DI values. The mean DI value of the control group was found to be 0.1573. When considered together with patient associated DI measurements, the control group distribution provides a baseline for discussing whether the observed DI values are consistent with the expected deformability range or tend to shift toward altered mechanical behavior. Therefore, this control data set serves as a significant comparative reference for the interpretation of remission compatibility and possible deviations in deformability behavior.

**Figure 4.**
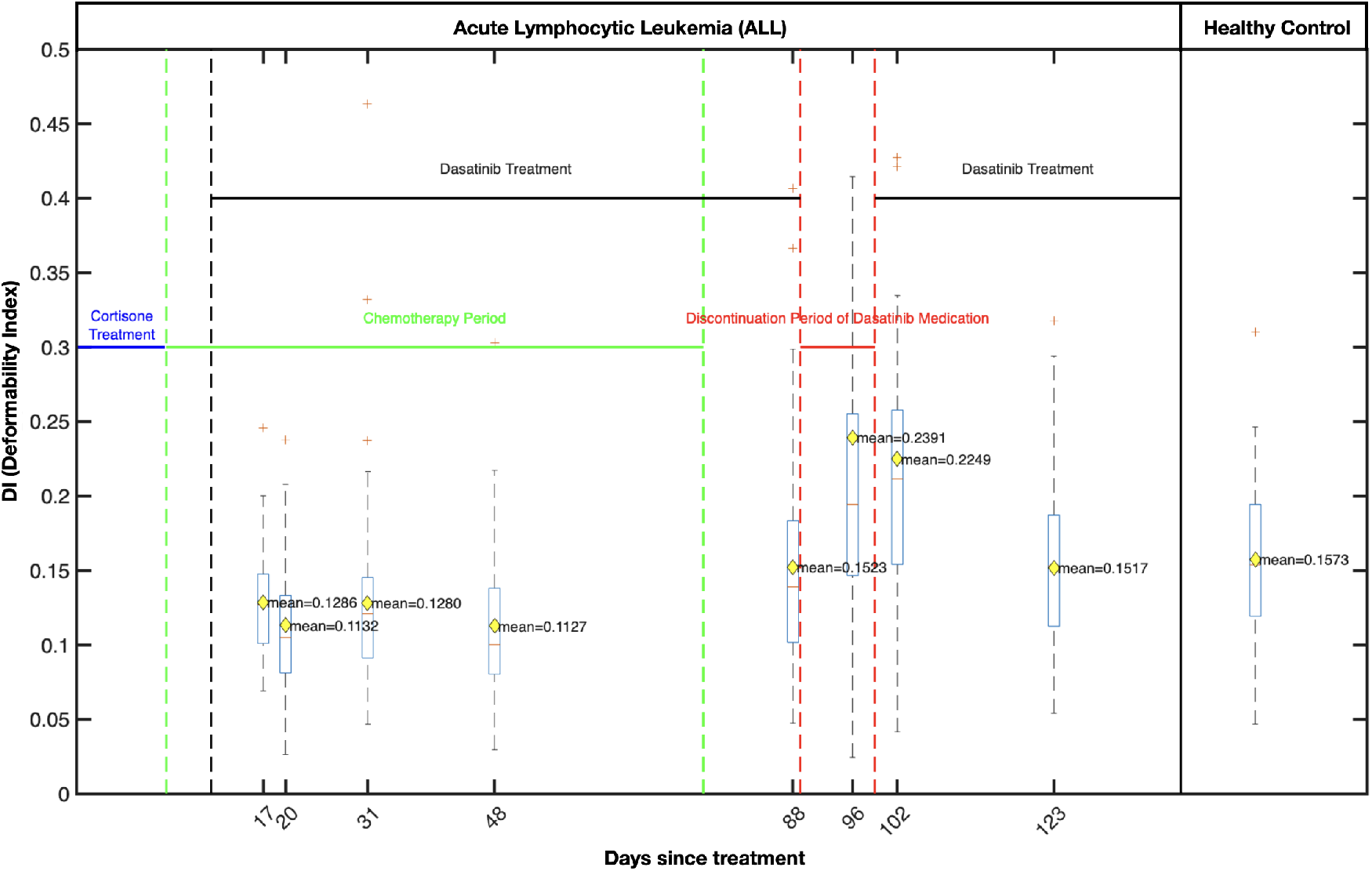
Boxplot of the DI measured at different time points during the treatment course. The figure illustrates changes in RBC deformability across the corticosteroid treatment, chemotherapy period, dasatinib treatment and discontinuation phases. Dashed vertical lines indicate transitions between treatment stages. Rightmost measurement is the DI value obtained from the control group which yielded a mean value of DI_mean_ = 0.1573.

As illustrated in Fig.5a, pairwise t-tests were performed between the DI measurements of the samples, in order to further assess whether the observed differences between measurement time points were statistically significant. The analysis indicated that several comparisons between earlier and later treatment stage measurements were statistically significant, supporting the variation observed in the data, shown in Fig.5b. In particular, the June and July samples indicated that there were no significant differences, which supports the differentiation of the chemotherapy region groupwise. Similarly, the August 12 sample showed a significant difference for August 20 and August 26 since these two samples were taken while the patient discontinued his dasatinib medication. Lastly, the September 16 sample was found to have significant differences for all samples except the August 12 sample, which is expected regarding the medication condition of the patient. These findings support the interpretation that RBC deformability changed non-uniformly throughout the treatment course.

**Figure 5.**
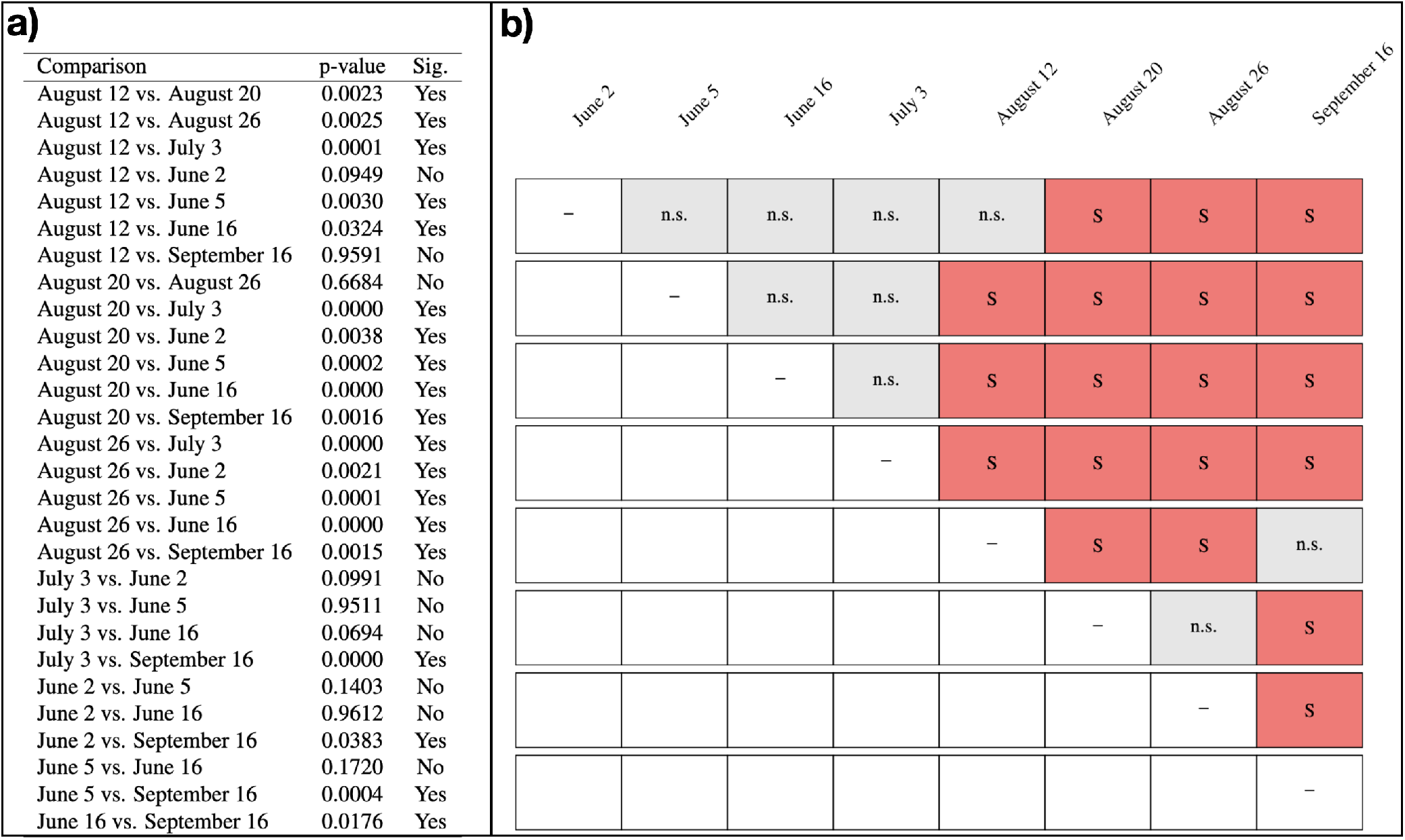
a) Pairwise t-test results for DI measurements between different sampling dates. Comparisons including the control sample are not shown. b) Pairwise comparison matrix of DI measurements across sampling dates based on t-test results. Dates are ordered chronologically from June 2 to September 16. Red boxes indicate statistically significant differences, whereas gray boxes indicate non-significant comparisons.

Furthermore, throughout the experiment, hemogram data were also obtained from the hospital in order to analyze the correlation of each hemogram element with respect to the DI measurements, as given in Fig.6a. The inferred Pearson correlation coefficients were then used to create a correlation heatmap, as illustrated in Fig.6b. The heatmap provided a compact visual summary of the relative strength and direction of these correlations in the hematological parameters measured.

**Figure 6.**
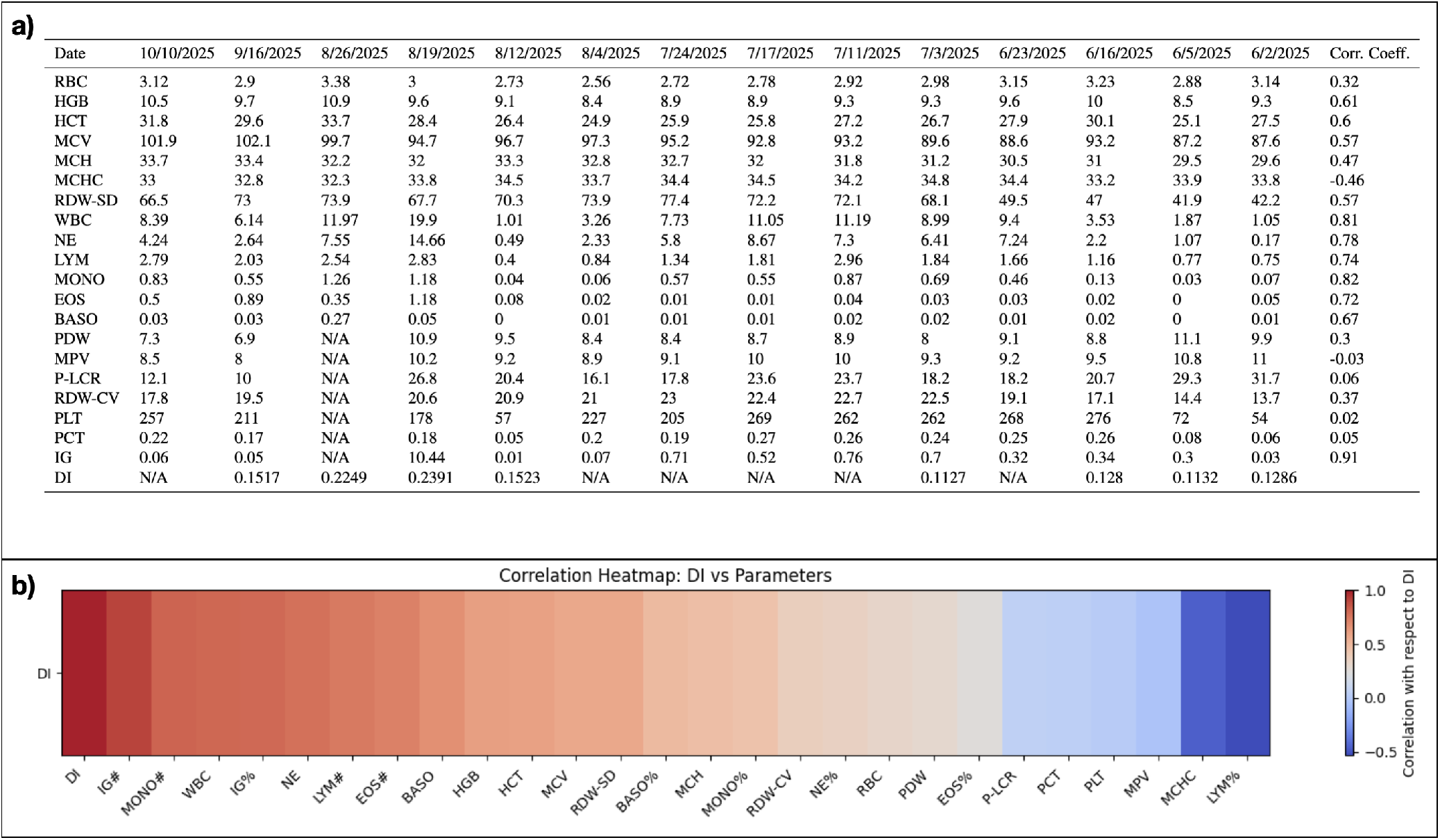
a) Hemogram test belongs to the patient. Each CBC (complete blood count) variables correlation coefficients with respect to DI was given in the last column. b) Correlation heatmap showing Pearson correlation coefficients between DI and hematological parameters.

Moreover, a principal component analysis (PCA) was performed in order to investigate how the measured DI values are distributed in relation to few important parameters within the multivariate space of the hemogram. The aim of this analysis was to assess whether DI follows a pattern of variation similar to erythrocyte, leukocyte, or platelet related hematological variables, and to visualize possible directional tendencies and groupings among these parameters. The PCA biplot using selected hemogram variables is presented in Fig.7, while the supplementary PCA biplot that includes all complete parameters is shown in Fig.8. The resulting distribution showed that DI is positioned closer to variables that are related to the leukocyte while it appears relatively separated from the direction of the parameters of the erythrocyte and platelet.

**Figure 7.**
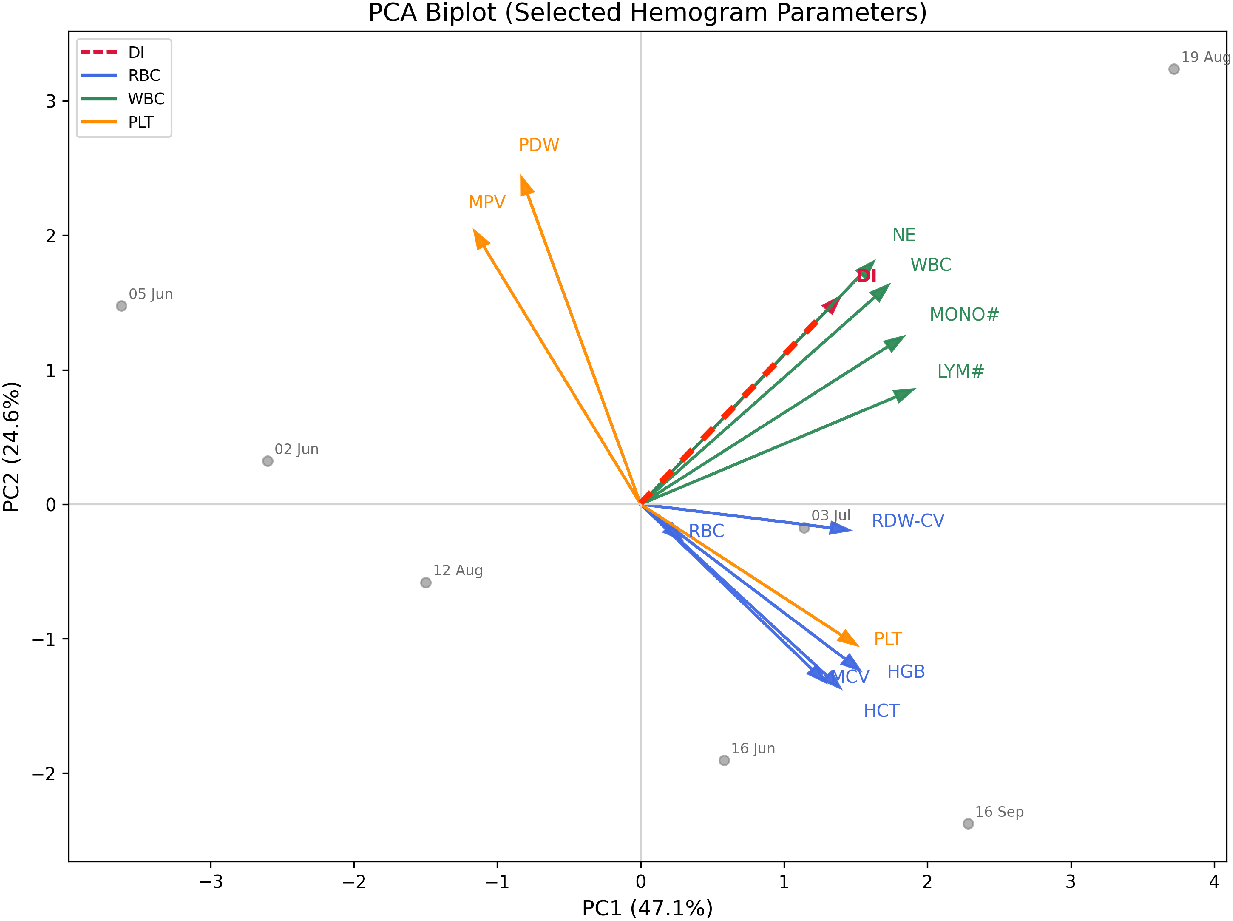
PCA biplot of selected hemogram parameters, including DI, RBC-related variables (RBC, HGB, HCT, MCV, RDW-CV), WBC-related variables (WBC, NE, LYM#, MONO#), and platelet-related variables (PLT, MPV, PDW). Sample dates are shown as gray points, while arrows represent variable loadings in the PC1–PC2 space after z-score standardization. PC1 and PC2 explained 47.1 % and 24.6% of the total variance, respectively. DI was observed to be positioned closer to leukocyte-related variables, particularly WBC and NE.

**Figure 8.**
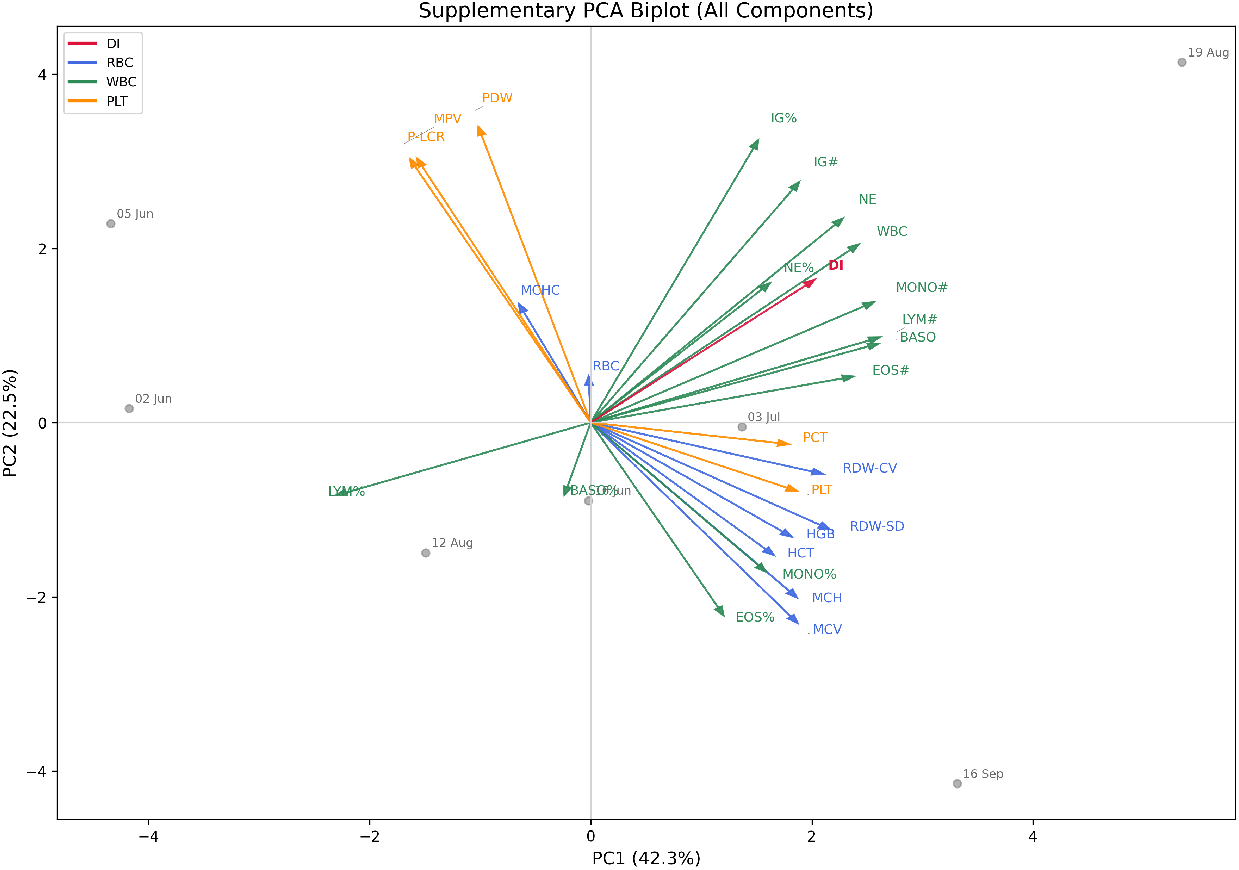
PCA biplot including all complete hemogram components together with DI. PC1 and PC2 explained 42.3% and 22.5% of the total variance, respectively. The broader analysis supported the main pattern observed in the reduced model, with DI aligning more closely with leukocyte-associated variables.

## Discussion

The present study investigated whether RBC deformability measured by optical tweezers could provide complementary information during the treatment course of an patient diagnosed with ALL. The results demonstrated that the deformability index was not constant throughout the treatment period but varied across different stages. In particular, relatively low DI values were observed during the early phase of chemotherapy, followed by a gradual increase during subsequent measurements.

These findings suggest that RBC deformability may reflect not only the intrinsic mechanical properties of erythrocytes but also broader systemic alterations associated with leukemia and its treatment.^28^ In ALL, leukemic infiltration of the bone marrow disrupts normal hematopoiesis and contributes to anemia.^9,29^ Accumulation of lymphoblasts within the marrow suppresses erythropoietic precursor cells and alters the bone marrow microenvironment, thereby affecting both red cell production and survival.^9,30^ These changes are further associated with alterations in stromal cell function and erythropoietic regulatory mechanisms.^31^

Beyond impaired erythropoiesis, systemic inflammation and oxidative stress associated with leukemia may compromise RBC membrane integrity and cytoskeletal organization, ultimately influencing erythrocyte mechanical behavior.^32^ This interpretation is consistent with previous reports describing structural abnormalities of RBCs in leukemic patients and altered blood rheology in leukemia.^22,23^ Treatment-related effects may also contribute to these changes. In vitro studies have demonstrated that several anticancer agents, including paclitaxel, carboplatin, doxorubicin, and cyclophosphamide, can modify RBC biophysical properties, suggesting that chemotherapy exposure itself may affect erythrocyte mechanics.^33^

The temporal evolution of DI values observed in this study is particularly noteworthy. According to Fig. 4, the relatively low deformability values measured during the early treatment phase may be associated with disease burden and chemotherapy exposure, whereas the subsequent increase in DI could indicate partial recovery of RBC mechanical properties during clinical improvement and remission. Similar trends have been reported during hematopoietic stem cell transplantation, where erythrocyte elongation index gradually increased between days 5 and 30 after transplantation, suggesting that RBC deformability may reflect changes occurring within the bone marrow microenvironment.^34^

Nevertheless, these findings should be interpreted cautiously. The present study does not permit the direct separation of the individual contributions of leukemia progression, systemic inflammation, and treatment-related effects. Therefore, the observed changes should be regarded as an association between the treatment course and the mechanical response of RBCs rather than evidence of a direct causal relationship.

In addition, the relatively long lifespan of RBCs, approximately 120 days,^35,36^ should be taken into account when interpreting deformability measurements. Consequently, the measured DI values likely represent a heterogeneous population of erythrocytes produced under different physiological conditions and treatment stages. This complexity should be considered when relating mechanical properties to specific disease states or therapeutic interventions.

Interestingly, the later measurements exhibited DI values that approached the average deformability levels observed in healthy controls. In particular, comparison with the control distribution revealed that the remission-phase measurement was among the patient values showing the greatest overlap with the healthy reference range. Although based on a single case, this observation raises the possibility that remission may be accompanied not only by improvements in conventional hematological parameters but also by a partial restoration of RBC mechanical properties. Such findings suggest that erythrocyte deformability may provide complementary information regarding physiological recovery beyond standard laboratory measurements.

Correlation analysis and principal component analysis provided additional insights into the relationship between RBC deformability and hematological variables. Interestingly, DI appeared to align more closely with leukocyte-related parameters than with the principal erythrocyte-related axis. Although these multivariate associations should be interpreted cautiously, they suggest that RBC deformability may reflect broader systemic and hematological changes occurring during treatment rather than representing an isolated erythrocyte characteristic. Therefore, alterations in RBC mechanics may serve as an integrated indicator of the overall biological state of the patient.

Several limitations of the present study should be acknowledged. First, the findings are based on a single patient and a limited number of measurement time points, preventing generalization to the broader ALL population. Second, the changing treatment conditions and the observational nature of the study make it impossible to distinguish the individual contributions of disease progression, inflammation, and chemotherapy exposure. Finally, the statistical analyses, including correlation analysis and PCA, should be regarded as exploratory because of the limited sample size. Consequently, the present findings should be considered preliminary evidence rather than definitive proof and warrant validation in larger longitudinal studies involving multiple patients.

## Conclusion

This study demonstrates that RBC deformability measured by optical tweezers changes dynamically throughout the treatment course of an elderly patient with ALL. Relatively reduced DI values were observed during the early stages of treatment, whereas higher values were detected in later measurements, suggesting that erythrocyte mechanical properties may vary in parallel with disease status and treatment-associated physiological changes.

These findings support the potential utility of optical tweezers-based RBC deformability analysis as a complementary biophysical approach for monitoring leukemia. Although the present observations are based on a single patient and should be considered preliminary, they provide a rationale for larger longitudinal studies aimed at evaluating whether RBC deformability can serve as a reproducible indicator of disease activity, treatment response, and remission in ALL.

## Acknowledgements and Fundings

This study was supported by a grant from the Presidency of the Republic of Türkiye, Presidency of Strategy and Budget (SBB) (Grant No. 2026K12-283013) and Scientific and Technological Research Council of Türkiye (TUBITAK, Grant No:124F223).

## Author contributions statement

T.P. and B.A. conceived the experiment, T.P. and Y.K. conducted the experiments, N.A. and B.B.B. supplied the sample. T.P., Y.K., B.B.B., N.A. and B.A. analysed the results. All authors reviewed the manuscript.

## Notes

### Competing Interest Statement

The authors have declared no competing interest.

